# Anti-NMDA receptor encephalitis in mice induced by active immunization with conformationally-stabilized holoreceptors

**DOI:** 10.1101/467902

**Authors:** Brian E. Jones, Kenneth R Tovar, April Goehring, Nana J. Okada, Eric Gouaux, Gary L. Westbrook

**Author notes:** **To whom correspondence should be addressed**: Gary L. Westbrook, Vollum Institute, L474, Oregon Health and Science University 3181 SW Sam Jackson Park Road Portland OR, 97239. current address: Neuroscience Research Institute, University of California, Santa Barbara, Santa Barbara, CA 93016. current address: Department of Psychology, 16 Barker Hall, University of California, Berkeley, Berkeley, CA 94720.

## Abstract

Autoimmunity to membrane proteins in the central nervous system has been increasingly recognized as a cause of neuropsychiatric disease. A key recent development was the discovery of antibodies to NMDA receptors in limbic encephalitis, characterized by cognitive changes, memory loss, seizures and sometimes long-term morbidity or mortality. Treatment approaches and experimental studies have largely focused on the pathogenic role of these autoantibodies. Passive antibody transfer to mice has provided useful insights, but does not produce the full spectrum of the human disease. Here we describe a *de novo* autoimmune mouse model of anti-NMDA receptor encephalitis. Active immunization of immune competent mice with conformationally-stabilized, native-like NMDA receptors induced a fulminant encephalitis that was strikingly similar to the behavioral and pathologic characteristics of human cases. Our results provide evidence of neuroinflammation and immune cell infiltration as early and robust features of the autoimmune response. Use of transgenic mice indicated that mature T cells as well as antibody-producing cells were required for disease induction. Our results provide new insights into disease pathogenesis as well as a platform for testing mechanisms of disease initiation and therapeutic approaches.

**One Sentence Summary:** We report an active immunization model of anti-NMDA receptor encephalitis in mice that recapitulates the features of the clinical disease, provides new insights into the pathophysiology, and offers a platform for investigation of new therapeutic interventions.

## Introduction

Limbic encephalitis, characterized clinically by behavioral changes, psychosis, memory impairment, and seizures is associated with hyperintense MRI signal in limbic structures *(1–3)*. In some cases, this discrete clinical syndrome has been attributed to Herpes simplex virus, but the underlying cause in other cases has remained unknown. In 2007, autoantibodies targeting *N*-methyl-D-aspartate (NMDA) receptor were discovered in a subset of these patients *(4, 5)*. With recognition of anti-NMDA receptor encephalitis as a clinical syndrome, diagnostic tests have revealed that the disease is surprisingly common in patients of all ages. Initially considered as one of the paraneoplastic disorders, it is now clear that most patients do not have detectable tumors *(6, 7)*, underscoring our lack of understanding of the underlying etiology. Patient-derived samples suggest a role for antibodies directed at NMDA receptor subunits in the pathogenesis, leading to treatments to reduce antibody titers with plasmapheresis or immunosuppression *(8)*. However, recovery following standard treatments can be prolonged and incomplete *(7, 9–11)*.

Passive transfer of antibodies from affected patients to mice have suggested that NMDA receptor antibodies can cause hypofunction in NMDA receptor-mediated synaptic transmission *(6, 12–14)*. Yet, the onset of symptoms in patients is usually insidious making it difficult to determine the initiating immunological factors, or the relative roles of the characteristic neuroinflammation and receptor hypofunction in the disease process *(12, 15–17)*. A robust animal model of the disease has the potential to address such issues.

To develop a mouse model of autoimmune encephalitis, we postulated that immunization with fully assembled receptors could be important in triggering the disease. Thus we used active immunization with intact native-like NMDA receptors composed of GluN1-GluN2B tetramers embedded in liposomes. Subcutaneous injection of these NMDA receptor proteoliposomes induced fulminant encephalitis within 4 weeks in young adult mice. The mice demonstrated behavioral changes, seizures as well as histological features of neuroinflammation and immune cell infiltration that were most prominent in the hippocampus. The presence of NMDA receptor antibodies was confirmed by immunohistochemistry and western blot. CD4^+^ T cell infiltration was a robust early feature, and both mature T and B cells were required for disease induction.

## Results

### Active immunization with NMDA receptor holoprotein induced encephalitis

We used a purified GluN1/GluN2B NMDA fully-assembled tetrameric receptors (holoreceptors) as described in methods to immunize C57Bl6 adult mice (Fig. 1A). Subcutaneous immunization with NMDA receptors in proteoliposomes was followed by a booster two weeks later. Littermate control cohorts were injected only with liposomes or with saline. Symptoms began to appear by four weeks, and by six weeks post-immunization nearly all of the proteoliposome-treated mice (86%) exhibited abnormal home cage behavior (Fig. 1B), indicating the high incidence of symptoms following holoprotein immunization. The most prominent behavioral changes were hyperactivity (86%) followed by tight circling (50%), overt seizures (21%) and hunched back/lethargy (11%), (Fig. 1C & Movie S1-S5). Cumulative symptom scores (described in methods) were significantly higher in proteoliposome-treated mice (proteoliposome v. liposome p < 0.0001, proteoliposome v. saline p < 0.0001, liposome v. saline p < 0.9999; Kruskal-Wallis, Dunn’s multiple comparisons post hoc; n=28/treatment group). As the presence of symptoms was assessed with two-minute daily observation periods, it is likely that seizure prevalence in particular was underestimated. Proteoliposome-treated mice also had increased mortality by six weeks post-immunization (n=8/56 proteoliposome, n=0/56 controls; p < 0.0005; Log-rank test).

**Figure 1.**
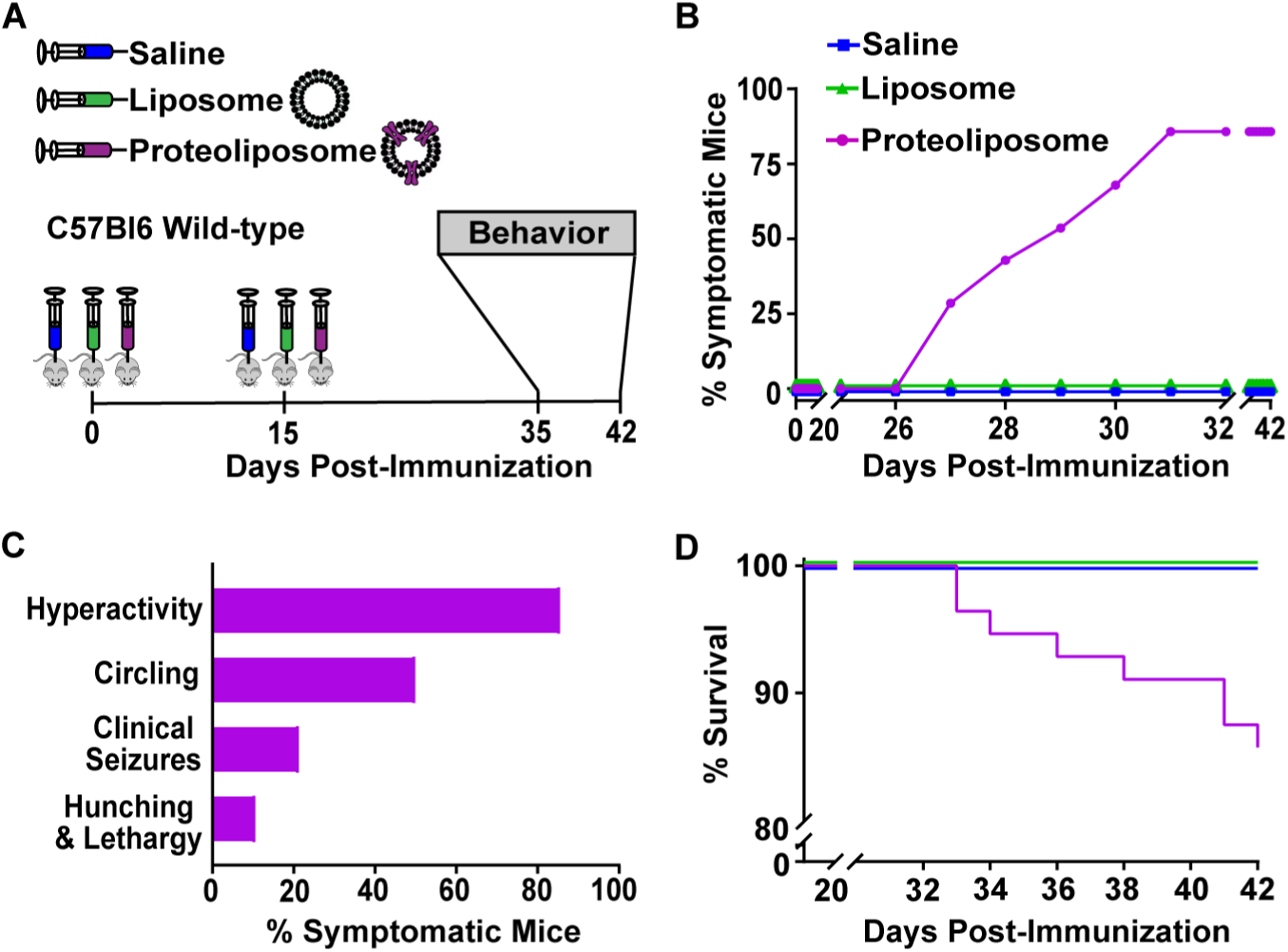
Proteoliposome-treated mice show a pronounced phenotype. (A) Timeline of treatment and behavioral testing. Adult wild-type received subcutaneous injections at day 0 and at day 15 with proteoliposome (purple) or control - liposome (green) or saline (blue). (B) Symptoms were plotted from the first injection. Only proteoliposome-treated mice displayed clinical signs by six weeks post-immunization. (C) Symptoms in proteoliposome-treated mice included locomotor hyperactivity and circling as well seizures and abnormal posture. (D) Kaplan-Meier survival plot revealed significant mortality in proteoliposome treated-mice.

To assess behavioral phenotype, we used a battery of standardized tests. Open field testing (Fig. 2A) confirmed a hyperactive locomotor phenotype with nearly double the distance traveled in proteoliposome-treated mice (proteoliposome. 5472 ± 525.4 cm, n =26; liposome = 3207 ± 111 cm, n = 28; saline = 3093 ± 84.9 cm; n=28; proteoliposome v. liposome p = 0.0002; proteoliposome v. saline p < 0.0001; liposome v. saline p > 0.9999; Kruskal-Wallis test with Dunn’s multiple comparisons post hoc). Proteoliposome-treated mice also showed increased variability in the open ranging from near immobility to extreme hyperactivity. Nest building, indicative of complex stereotyped behavior, was severely compromised in the proteoliposome-treated mice (Fig. 2B). At six weeks post-immunization, control mice created precise nests whereas proteoliposome-treated mice barely disturbed the nestlets as indicated by a lower nesting score: proteoliposome (24,48 hours) = 1 ± 0.29 and 1.48 ± 0.39, n=27; liposome = 4.60 ± 0.14, and 4.78 ± 0.11, n = 28; saline = 4.21 ± 0.14 and 4.75 ± 0.12, n = 28. proteoliposome v. liposome p < 0.0001; proteoliposome v. saline p < 0.0001; liposome v. saline p = 0.3603 (24 hours); proteoliposome v. liposome. p < 0.0001; proteoliposome v. saline p < 0.0001; liposome v. saline p > 0.9999 (48 hours); Kruskal-Wallis with Dunn’s multiple comparisons post hoc). In the zero maze, often used as a measure of anxiety-like behavior, proteoliposome-treated mice spent more time in the normally aversive open-arm (% time open-arm: proteoliposome = 27.90 ± 3.70, n = 18; liposome = 16.08 ± 0.71, n = 28; saline = 16.32 ± 0.96, n = 28; proteoliposome v. liposome. p = 0.0065; proteoliposome v. saline. p = 0.0081; liposome. v. saline. p > 0.9999; Kruskal-Wallis with Dunn’s multiple comparisons post hoc). There was no statistical difference in the total distance moved in the open-arm indicating that hyperactivity could not explain the observed phenotype (proteoliposome: 175.4 ± 26.18 cm; liposome:118.7 ± 9.41 cm; saline; 130.4 ± 15.67 cm; proteoliposome v. liposome p = 0.1472; proteoliposome v. saline. p = 0.2280; liposome. v. saline. p = 0.6489; 2-way ANOVA with Holm-Sidak’s multiple comparison). Although proteoliposome-treated mice were marginally more active compared to liposome controls in the closed arm, this difference was not supported by comparing proteoliposome with saline controls (proteoliposome v. liposome p = 0.0239; proteoliposome v. saline. p = 0.1836; liposome. v. saline. p = 0.2583; 2way ANOVA with Holm-Sidak’s multiple comparison test). These results indicated that immunization with NMDA receptor holoprotein induces a striking behavioral phenotype consistent with encephalitis.

**Figure 2.**
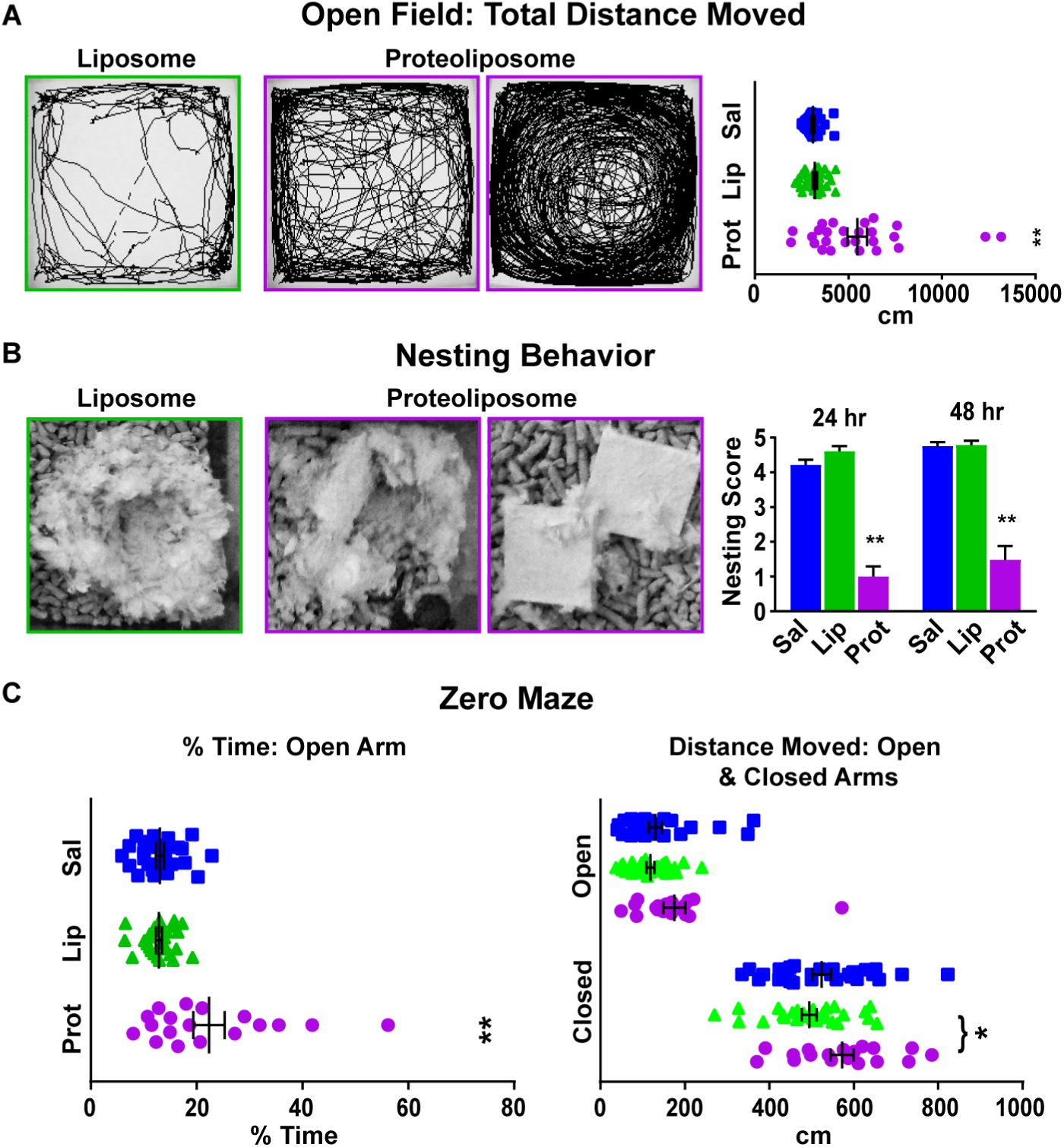
Behavioral assessment. (A) Representative movement traces in the open field from a control mouse (field outlined in green) and two proteoliposome treated-mice (field outlined in purple) illustrate the hyperactivity phenotype. Total distance moved in the treatment groups are plotted at right: proteoliposome (purple), liposome (green) and saline (blue) purple. (B) Representative images of a normal nest created by a control mouse compared to the partial or incomplete nests in two proteoliposome treated-mice. Nests were scored at 24 and 48 hours and quantified as shown at right. (C) Mice were assessed in the zero for time spent in the open arm (left) and for the total distance moved in the open and closed arms.

### Neuroinflammation and peripheral leukocyte infiltration

Autoimmune encephalitis is associated with characteristic histopathology including cell infiltrates as well as inflammation and neuronal loss *(15–18)*. In proteoliposome-treated mice, hematoxylin-labeled perivascular cuffing was prominent in multiple CNS regions including the hippocampus and neocortex (Fig. 3A, right panels). Patchy areas of cell death were also present (Fig. 3B, right panels). No signs of inflammation or cell death were present in controls (Fig. 3A, B, left panels). Immunolabeling with GFAP and Iba1 revealed a pronounced inflammatory response in proteoliposome-treated mice (Fig. 4A,B, right panels). Total GFAP immunoreactivity in the hippocampus was much higher than controls (proteoliposome: 42.5 × 10^8^ ± 8.80 × 10^8^; control: 6.33 × 10^8^ ± 1.64 × 10^8^; proteoliposome v. control. p = 0.0022; Mann Whitney test; n=6 mice /group). Iba1 labeling revealed foci of microgliosis compared to scattered diffuse labeling in controls (Fig. 4B). Total Iba1 immunoreactivity in the hippocampus was 60.9 × 10^8^ ± 14.3 × 10^8^; in proteoliposome-treated mice compared to 28.5 × 10^8^ ± 1.64 × 10^8^ in controls: 28.5 × 10^8^ ± 1.64 × 10^8^ (p = 0.0022, Mann Whitney test; n=6 / group).

**Figure 3.**
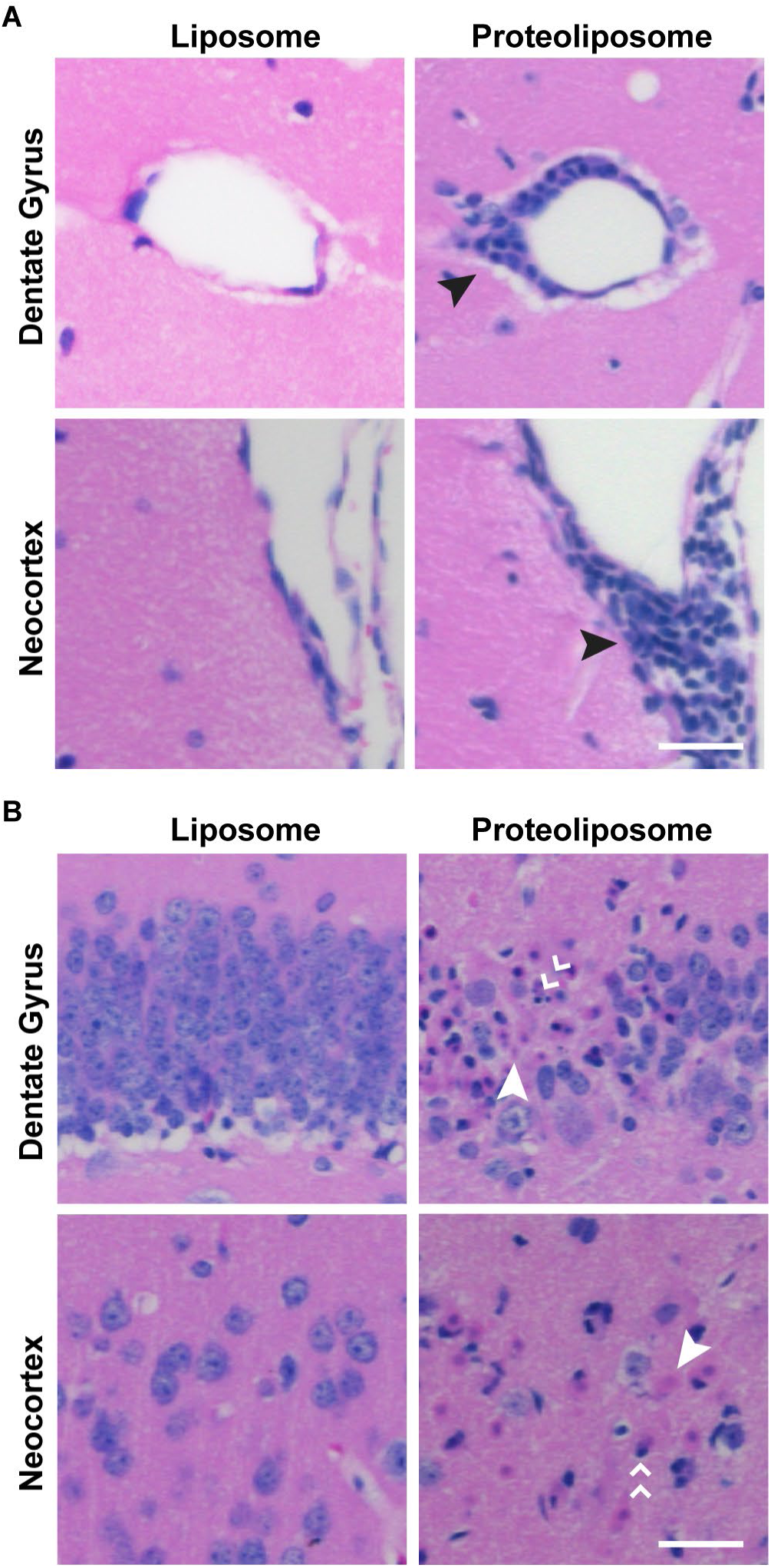
Histological evidence of CNS inflammation. (A) Representative Images of perivascular cuffing (black arrow, right panels) in hippocampal and neocortical tissue from a proteoliposome-treated mouse. Matched samples from liposome-treated controls are shown at left. (B) Representative tissue sections also showed discrete areas of cell loss including karyolysis (single white arrow) and pyknosis (double white arrow) in a proteoliposome-treated mouse (right), but not in a control (left). Scale bar: 100μm.

**Figure 4.**
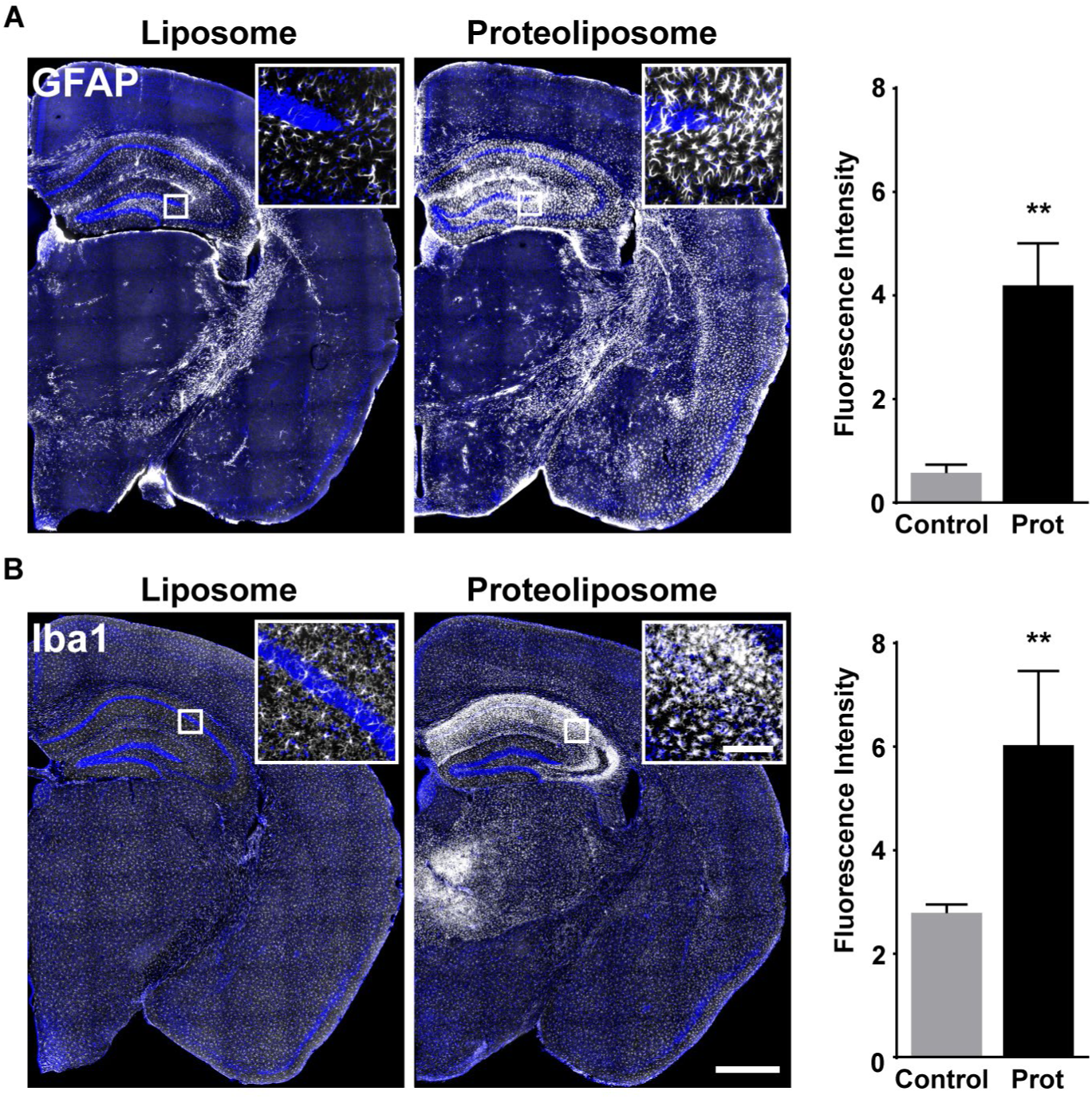
Evidence for reactive gliosis. (A) Immunofluorescence for GFAP (white) was markedly enhanced in proteoliposome-treated mice (right) compared to controls (left). The insets are higher magnification images from the hippocampus to show individual astrocytes. The histogram shows quantification of GFAP immunofluorescence (see methods). (B) Labeling with the microglial marker Iba1 (white) was also greatly increased in proteoliposome-treated mice (right) compared to control (left), particularly in the hippocampus and thalamus. Higher magnification inset show intense microglia in the hippocampus of proteoliposome-treated mouse compared to the scattered microglial labeling in control. Immunofluorescence was quantified in the histogram as for GFAP. Scale bar: 1000μm (inset 100μm).

There was also pronounced CNS infiltration by peripheral immune cells at six weeks post-immunization. Labeling with the pan-leukocyte marker CD45R was robust in the hippocampus (Hipp), striatum (Str), thalamus (Thal), amygdala (Amyg) and neocortex (Ctx), (Fig. 5A, right panel). Control mice showed sparse CD45R labeling (Fig. 5A, left panel). Mean CD45R^+^ cell densities for control- and proteoliposome-treated mice are summarized in Table 1. Peripheral immune cells including activated macrophages/microglia (Galectin3^+^), plasma cells (CD138^+^), helper T cells (CD4^+^), B cells (CD20^+^) as well as cytotoxic T cells (CD8^+^) also were strikingly increased in the brains of proteoliposome-treated mice (Fig. 5B, right panel). In contrast liposome- or saline-treated mice showed sparse or absent immunoreactivity (Fig. 5B, left panel). Quantification of immune cell subtypes in the hippocampus is summarized in Table 1. Although mice at 3 weeks post-immunization did not show prominent clinical features, neuroinflammation and immune cell infiltration were already present at this early time point (Table 1 and Fig. S2).

**Figure 5.**
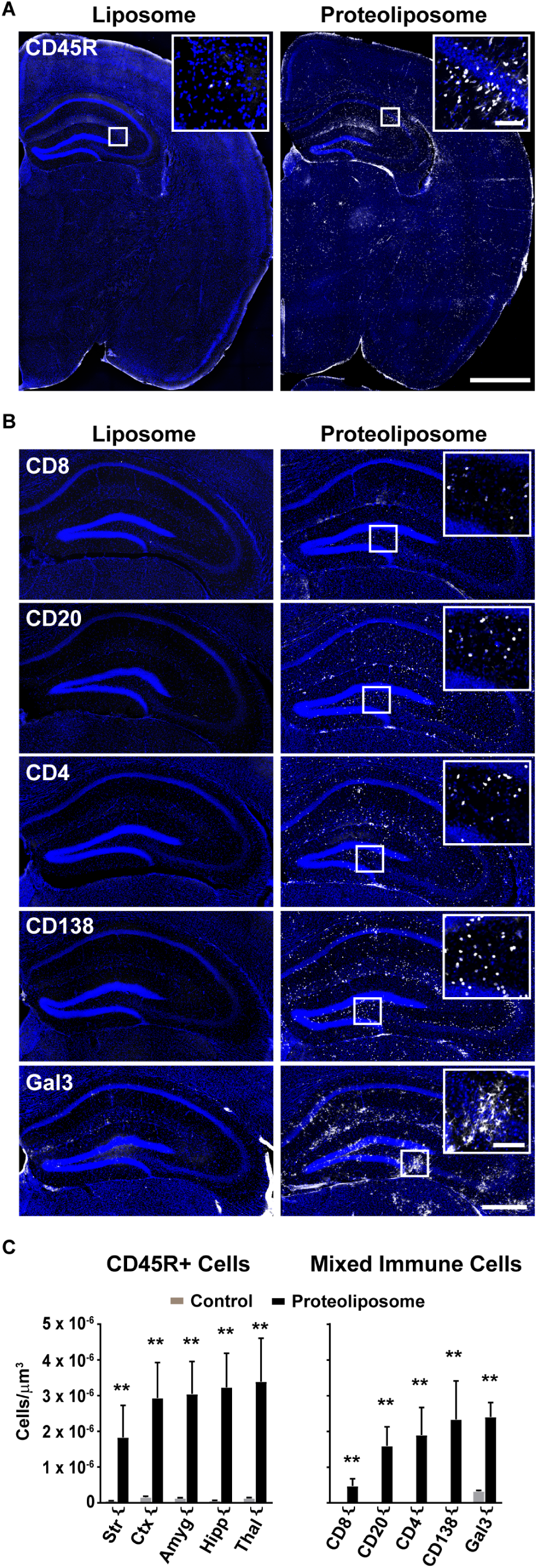
Prominent immune cell infiltration. (A) Robust expression of the panleukocyte marker, CD45R in a proteoliposome treated-mouse (right). Insets show labeling of individual CD45R^+^ cells (white) from the indicated region of the hippocampus. Coronal sections from control mice had few if any CD45R^+^ cells (left). Scale bar: 1000μm (inset 100μm). (B) Control mice (left) had sparse or absent labeling for a battery of immune cell markers (CD8^+^, CD4^+^, CD20^+^, CD138^+^, Gal3^+^). Proteoliposome treated-mice showed significant increases in immune cell infiltrates as indicated with the insets from the hippocampal region; [Scale bar: 500μm (inset 200μm). (C) Left: CD45R^+^ density (cells/μm^3^) in striatum (Str), cortex (Ctx), amygdala (Amyg), hippocampus (Hipp) and thalamus (Thal) in proteoliposome-treated (black) and control mice (gray). Right: CD8^+^, CD4^+^, CD20^+^, CD138^+^, Galectin3^+^ cell densities (cells/μm^3^) in the hippocampus of proteoliposome treated (black) and control mice (gray). Only Gal3^+^ cells were of sufficient density to be seen on the histogram in control mice.

**Table 1.** Quantification of immune cell infiltration. Mean CD45R^+^ cell densities across sampled anatomical regions in control and proteoliposome-treated mice at 6 and 3 weeks post-immunization (± SEM). Mean immune cell subtype densities in the hippocampus of control and proteoliposome treated mice at 6 and 3 weeks post-immunization (± SEM).

| Mean CD45R cell densities by anatomical region: 6 weeks post-immunization |  |  |
| --- | --- | --- |
| Anatomical Region | Control (cells/$\mu\text{m}^3$) | Proteoliposome (cells/$\mu\text{m}^3$) |
| Striatum | $0.57 \times 10^{-7} \pm 0.05 \times 10^{-7}$ | $18.3 \times 10^{-7} \pm 8.94 \times 10^{-7} **$ |
| Cortex | $1.54 \times 10^{-7} \pm 0.32 \times 10^{-7}$ | $29.3 \times 10^{-7} \pm 9.92 \times 10^{-7} **$ |
| Amygdala | $1.32 \times 10^{-7} \pm 0.13 \times 10^{-7}$ | $30.4 \times 10^{-7} \pm 9.07 \times 10^{-7} **$ |
| Hippocampus | $0.59 \times 10^{-7} \pm 0.15 \times 10^{-7}$ | $32.3 \times 10^{-7} \pm 9.50 \times 10^{-7} **$ |
| Thalamus | $1.30 \times 10^{-7} \pm 1.89 \times 10^{-7}$ | $33.9 \times 10^{-7} \pm 12.1 \times 10^{-7} **$ |
| **p = 0.0022, Mann Whitney test; n=6 mice/group |  |  |
| Mean CD45R cell densities by anatomical region: 3 weeks post-immunization |  |  |
| Anatomical Region | Control (cells/$\mu\text{m}^3$) | Proteoliposome (cells/$\mu\text{m}^3$) |
| Striatum | $1.16 \times 10^{-7} \pm 0.15 \times 10^{-7}$ | $108.0 \times 10^{-7} \pm 54.5 \times 10^{-7}$ |
| Cortex | $2.33 \times 10^{-7} \pm 0.44 \times 10^{-7}$ | $46.7 \times 10^{-7} \pm 16.9 \times 10^{-7}$ |
| Amygdala | $0.85 \times 10^{-7} \pm 0.19 \times 10^{-7}$ | $47.3 \times 10^{-7} \pm 34.4 \times 10^{-7}$ |
| Hippocampus | $0.97 \times 10^{-7} \pm 0.16 \times 10^{-7}$ | $88.1 \times 10^{-7} \pm 17.5 \times 10^{-7} *$ |
| Thalamus | $1.53 \times 10^{-7} \pm 0.36 \times 10^{-7}$ | $28.9 \times 10^{-7} \pm 13.7 \times 10^{-7}$ |
| *p = 0.0286, Mann Whitney test; n=5 mice/group |  |  |
| Mean Immune cell subtype densities in hippocampus: 6 weeks post-immunization |  |  |
| Immune Cell Subtype | Control (cells/$\mu\text{m}^3$) | Proteoliposome (cells/$\mu\text{m}^3$) |
| CD8 | $0.026 \times 10^{-7} \pm 0.014 \times 10^{-7}$ | $4.74 \times 10^{-7} \pm 2.00 \times 10^{-7} **$ |
| CD20 | $0 \times 10^0 \pm 0 \times 10^0$ | $15.9 \times 10^{-7} \pm 5.33 \times 10^{-7} **$ |
| CD4 | $0.007 \times 10^{-7} \pm 0.006 \times 10^{-7}$ | $18.9 \times 10^{-7} \pm 7.69 \times 10^{-7} **$ |
| CD138 | $0.007 \times 10^{-7} \pm 0.007 \times 10^{-7}$ | $23.3 \times 10^{-7} \pm 10.7 \times 10^{-7} **$ |
| Galectin3 | $3.21 \times 10^{-7} \pm 0.265 \times 10^{-7}$ | $24.0 \times 10^{-7} \pm 4.05 \times 10^{-7} **$ |
| **p = 0.0022, Mann Whitney test; n=6 mice/group |  |  |
| Mean Immune cell subtype densities in hippocampus: 3 weeks post-immunization |  |  |
| Immune Cell Subtype | Control (cells/$\mu\text{m}^3$) | Proteoliposome (cells/$\mu\text{m}^3$) |
| CD8 | $0.009 \times 10^{-7} \pm 0.006 \times 10^{-7}$ | $0.10 \times 10^{-7} \pm 0.03 \times 10^{-7}$ |
| CD20 | $0.03 \times 10^{-7} \pm 0.03 \times 10^{-7}$ | $13.2 \times 10^{-7} \pm 9.47 \times 10^{-7} *$ |
| CD4 | $0.38 \times 10^{-7} \pm 0.11 \times 10^{-7}$ | $35.9 \times 10^{-7} \pm 16.5 \times 10^{-7}$ |
| CD138 | $0.07 \times 10^{-7} \pm 0.04 \times 10^{-7}$ | $13.9 \times 10^{-7} \pm 11.4 \times 10^{-7}$ |
| Galectin3 | $2.81 \times 10^{-7} \pm 0.34 \times 10^{-7}$ | $39.8 \times 10^{-7} \pm 14.0 \times 10^{-7}$ |
| *p = 0.0476, Mann Whitney test; n=5 mice/group |  |  |

### Receptor autoantibodies in serum from proteoliposome-treated mice

The above results indicate that neuroinflammation and immune cell infiltrates were a prominent feature of NMDA receptor immunized mice. Staining for mouse IgG as a proxy for the presence of autoantibodies, essentially outlined the hippocampus (Fig. S1), consistent with patterns observed in anti-NMDA receptor encephalitis. Serum pooled from proteoliposome-treated mice showed bands on Western blots corresponding to purified recombinant rat and *Xenopus* GluN1 subunit protein as well as *Xenopus* GluN2B. Although epitopes in the GluN1 amino-terminal domain (ATD) have been identified in some human cases *(19, 20)*, the serum from proteoliposome-treated mice also labeled a *Xenopus* GluN1 subunit that lacked the ATD domain. Serum from liposome or saline-treated mice did not recognize NMDA receptor subunits (Fig. 6A, middle panel). Serum from proteoliposome-treated mice collected at three weeks post-immunization did not label NMDA receptor subunits bands, but labeling was present in cell-based assays (Fig. S3), perhaps indicating a lower antibody titer at this early time point.

**Figure 6.**
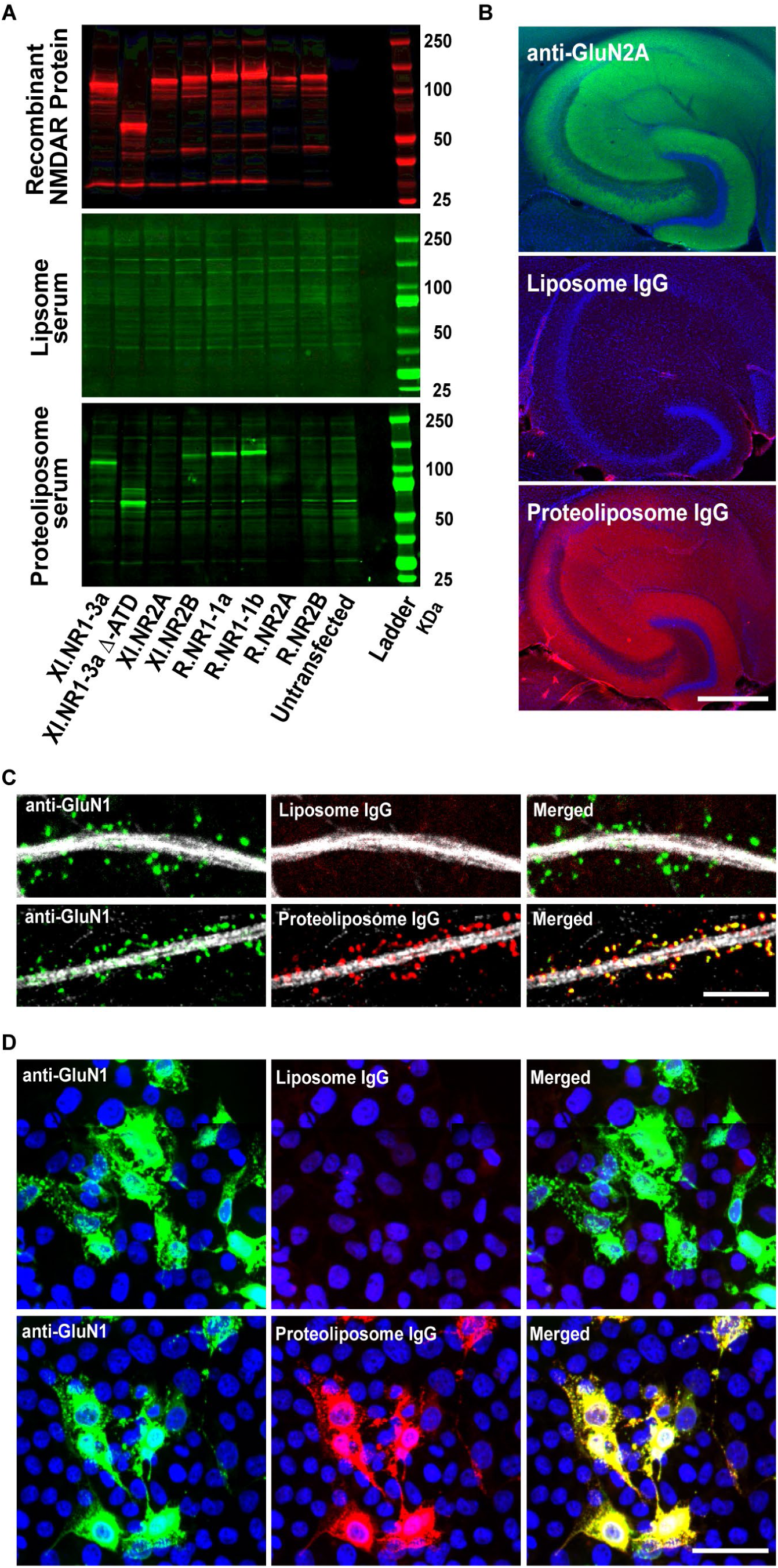
Detection of NMDA receptor subtype-specific serum antibodies and localization of IgG binding to NMDA receptors. (A) A battery of recombinant GFP-tagged NMDA receptor subunits from *Xenopus* (XI) and Rat (R) were blotted and detected with anti-GFP antibodies (red, top panel). Incubation with serum (1:100) derived from a liposome-treated mouse showed no specific labeling (green, middle panel). Serum (1:100) derived from a proteoliposome-treated mouse showed bands corresponding to GluN1 subunit isoforms in *Xenopus* and rat (Xl.GluN1-3a, R.GluN1-1a, R.GluN1-1b), *Xenopus* GluN2B as well as a *Xenopus* GluN1 lacking the ATD domain (XI.NR1-3a ΔATD). Blots are from representative liposome-treated and symptomatic proteoliposome-treated mice. (B) The pattern of immunoreactivity in hippocampal tissue sections for a commercial NMDA receptor antibody (GluN2A) in a wildtype mouse (left) matched that for purified IgG derived from proteoliposome-treated mice (right). IgG from liposome-treated mice had no labeling (middle). (Scale bar: 500μm). (C) Dendrites of cultured hippocampal neurons showed the expected punctate pattern of synapses using an anti-GluN1 antibody (green, left). IgG from proteoliposome-treated mice also showed punctate labeling along dendrites (bottom row, middle, red) that exactly matched anti-GluN1 labeling in the merge (bottom row, right, yellow). IgG from liposome-treated mice showed no labeling (top row, middle) although synapses were present as labeled with anti-GluN1 (top row, left and right, green).Dendritic shafts were labeled with anti-MAP2 antibody (gray) to enhance visualization, Scale bar: 5μm. (D) HEK293FT cells transfected with rat GluN1/2A subunits were labeled with anti-GluN1 antibody (left, top and bottom, green) as did IgG from proteoliposome-treated mice (bottom, middle, red). Merge panel shows colocalization (bottom, right, yellow). IgG derived from control mice (top middle panel, red) showed no labeling. Scale bar: 15μm.

Hippocampal labeling with purified IgG from proteoliposome-treated mice matched labeling with a GluN2A subunit-specific antibody (Fig. 6B, 6 weeks post-immunization). Purified IgG from controls showed no labeling. In cultured mouse hippocampal neurons (DIV14-21), only the IgG isolate from proteoliposome-treated mice co-localized with the GluN1 subunit specific antibody (Fig. 6C). The punctate labeling along dendrites and at dendritic spines follows the expected distribution of NMDA receptors at synapses. To provide a further assessment of subunit specificity in the IgG isolates, we used HEK293 cells to express and stain for NMDA receptor subunits. HEK cells were transfected with single subunit constructs or combinations of GluN1/2A and GluN1/2B. As observed in the primary hippocampal cultures, the proteoliposome IgG isolate co-localized with cells labeled with a GluN1 subunit-specific antibody (Fig. 6D). These data are consistent with Western blots and indicate the presence by 6 weeks post-immunization of polyclonal antibodies for GluN1 and GluN2B.

Serum from proteoliposome-treated mice did not acutely block NMDA receptor function, as assessed by whole-cell currents in cultured hippocampal neurons (Fig. 7A & B). NMDA (50 μM) was co-applied by local flow pipes either with serum from liposome-treated mice or serum from proteoliposome-treated mice (1:100 dilution). The NMDA-evoked current in the presence of serum from proteoliposome-treated mice was 95.9 ± 6.8% of that evoked by NMDA + serum from liposome-treated mice in the same neuron (n=8; p=0.23, paired t-test). In contrast, a 24-hour incubation with serum from proteoliposome-treated mice reduced synaptic activation of NMDA receptors. As shown in Figure 7C (top left), the slow components of EPSC barrages from neurons incubated in serum from liposome-treated mice were robust, but were greatly reduced by the NMDA receptor antagonist, D-AP5, as reflected by the rapid decay of the spontaneous EPSCs (Fig 7C, top right). After 24-hour incubation in serum from proteoliposome-treated mice, spontaneous EPSCs had reduced NMDA receptor mediated currents and thus were less sensitive to block by D-AP5 (Fig. 7C, bottom right). For neurons incubated in serum from liposome-treated mice total charge from spontaneous EPSCs was reduced to 44.1 ± 7.9% of control charge by D-AP5 (Fig. 7D). In contrast, D-AP5 reduced total charge to only 85.6 ± 6.0% of control charge in neurons incubated with serum from proteoliposome-treated mice (n=8 per group; p < 0.005, paired t-test, Fig. 7D), consistent with a reduction in NMDA receptor function at 24 hours.

**Figure 7.**
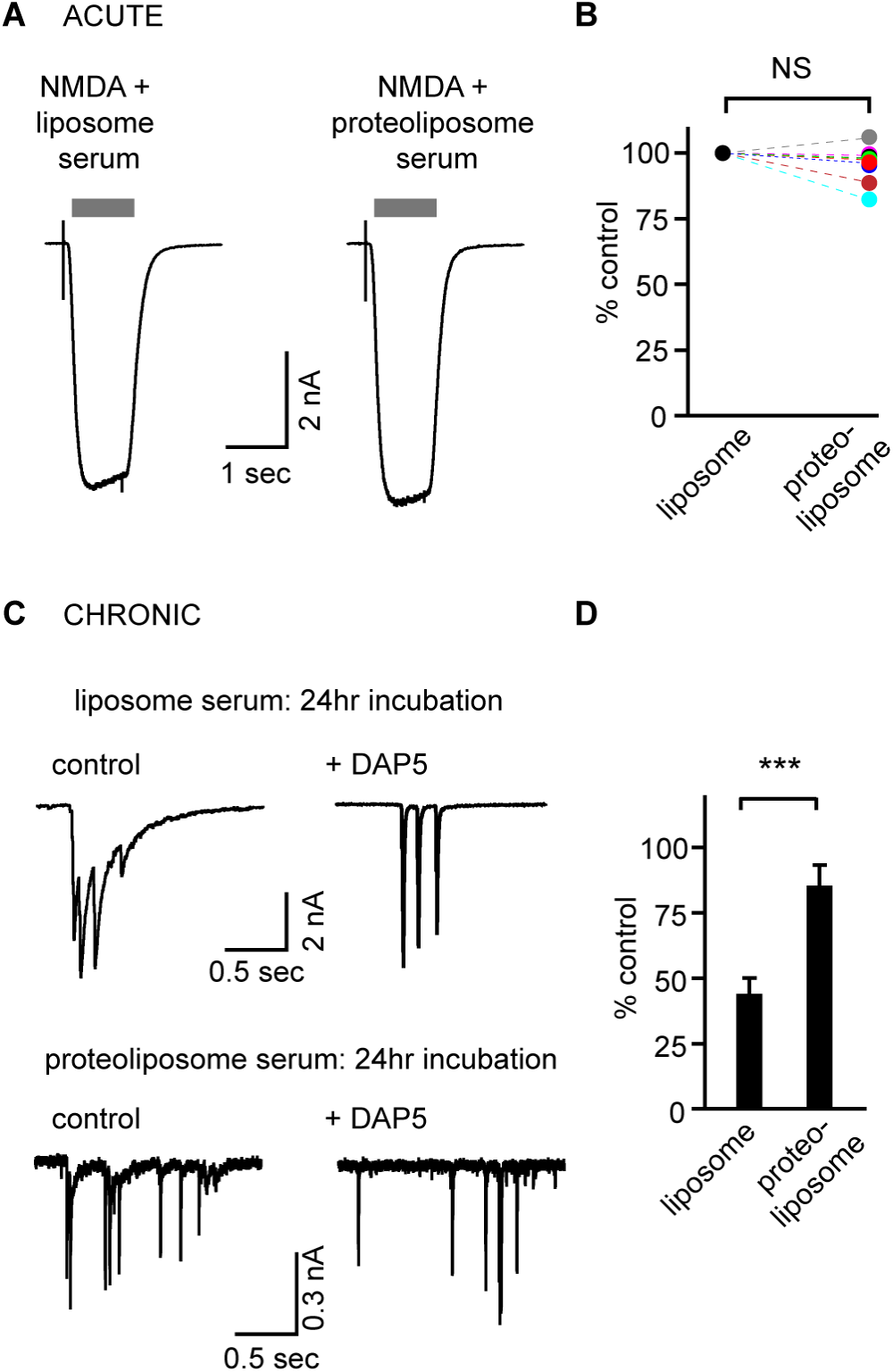
Serum-induced reduction in NMDA receptor function. (A) Representative whole-cell currents in a neuron evoked by acute flow-pipe application of NMDA + serum from liposome-treated, or NMDA + serum from proteoliposome-treated mice, respectively. Serum dilution was 1:100. (B) Evoked whole-cell NMDA currents for 8 neurons revealed no significant effect of proteoliposome serum on current amplitudes. (C, D) Spontaneous excitatory postsynaptic currents were recorded following 24-hour incubation (“chronic”) in serum from liposome or proteoliposome-treated mice. EPSCs in cells treated with serum from liposome-treated mice showed prolonged bursts of inward currents that were reduced by the NMDA receptor antagonist, D-AP5 (top left and right). The slow components of the NMDA receptor-mediated currents were less prominent after incubation with serum from proteoliposome-treated mice (bottom left) and were significantly less sensitive to block by D-AP5 (bottom right) as quantified by total charge transfer in the histogram in panel D.

### T cells as well as B cells are necessary for proteoliposome-induced encephalitis

Studies of anti-NMDA receptor encephalitis in human cases have focused on the role of antibodies *(4, 12, 13)*. Our results in proteoliposome-treated mice are consistent with the presence of B cell infiltrates as well as NMDA receptor autoantibodies. However, we saw both B and T cell infiltrates in proteoliposome-treated mice. To distinguish the roles of B and T cells in the mouse disease, we used two well-characterized mutant mouse lines that lack mature B cells or mature T cells *(21, 22)*. Consistent with a role for B cells in the pathophysiology, proteoliposome treatment of MuMt^−^ mice, which lack the capacity to generate an antigen specific antibody response, showed no behavioral or histological abnormalities at 6 and 12 weeks post-immunization. (Fig. S4). To evaluate the role of mature T cells we used Tcrα^−^ (“CD4, CD8 cell KO”) mice which lack mature helper T and cytotoxic T cells. All proteoliposome-treated Tcrα^−^ mice survived to 12 weeks without clinical signs of disease (Fig. 8A). Likewise Tcrα^−^ mice did not show histopathological evidence of gliosis or cell infiltrates and serum lacked detected anti-NMDA receptor antibodies (Fig. 8B-D). A cohort of immune competent wild-type mice immunized in parallel with the MuMt^−^ and Tcrα^−^ showed the expected phenotype, indicating the potency of the immunogen. These results confirm the expected role of B cells, but also indicate a requirement for mature T cells in disease pathogenesis.

**Figure 8.**
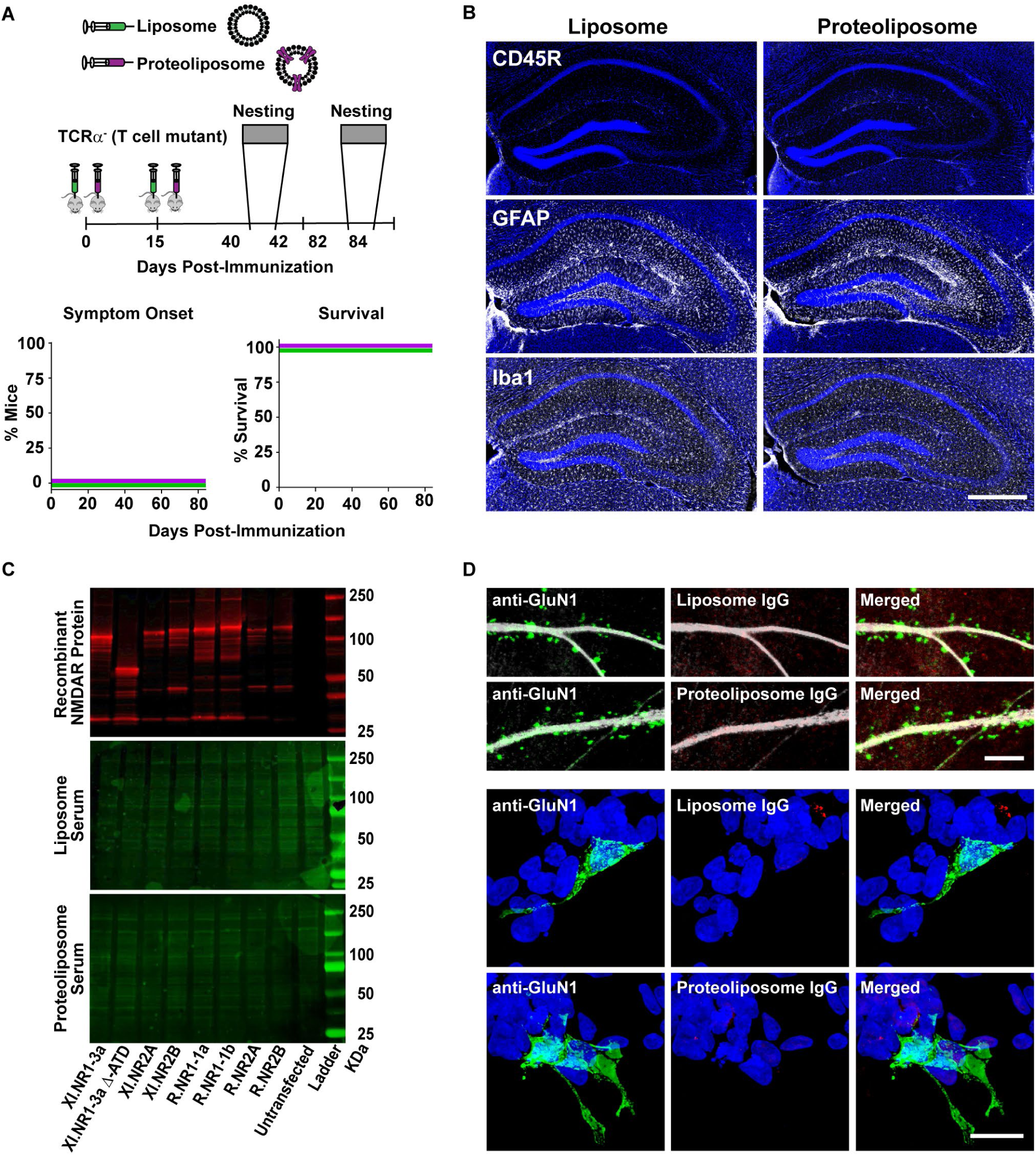
T cells are required to generate disease state in proteoliposome treated mice. (A) Tcrα^−^ mutant mice were immunized in parallel with a cohort of wildtype mice. None of the proteoliposome-treated mutant mice showed clinical or behavioral signs of encephalitis, or altered mortality rates by 12 weeks post-immunization (liposome = green; proteoliposome = purple). (B) Proteoliposome-treated TCRα^−^ mice did not show increased labeling with CD45R (top panels), GFAP (middle panels) or Iba1 (bottom panels) compared to liposome-treated TCRα^−^ mice. Scale bar: 500μm (C) A battery of recombinant GFP-tagged NMDA receptor subunits from *Xenopus* (XI) and Rat (R) were blotted and detected with anti-GFP antibodies (red, top panel). Incubation with serum (1:100) derived from a liposome- and proteoliposome-treated mouse showed no specific labeling (green, middle & bottom panel). (D) Purified IgG from proteoliposome-treated TCRα^−^ mice did not label synapses in mouse hippocampal neurons or HEK293FT cells expressing rat GluN1/2A subunits. The upper two rows show synapse with anti-GluN1 antibody (left, green) whereas IgG from liposome-treated mice (top middle, red) or IgG from proteoliposome treated-mice (bottom middle panel, red) showed no labeling. Scale bar = 5μm. The lower two rows show anti-GluN1 antibody labeling of HEK293FT cells expressing rat GluN1/2A subunits (left, green), but no labeling was detected using IgG from proteoliposome-treated or liposome-treated mice (bottom two rows, middle). Scale bar: 15μm.

## Discussion

The cause of the clinical syndrome of limbic encephalitis was an enigma for decades. The discovery of NMDA receptor antibodies in some of these patients was not only a surprise, but also opened the opportunity for better understanding of the causes and also treatments *(6)*. Animal models have played an important role in the study of neurological disorders including autoimmune diseases such as myasthenia gravis and multiple sclerosis *(23, 24)*. Key to the utility of an animal model is whether the model accurately mimics the essential features of the disease. For example, peptide fragments from myelin have long been used to generate experimental autoimmune encephalomyelitis (EAE) with its incumbent clinical symptoms to test the molecular basis and therapeutic approaches in MS *(23, 25, 26)*. Likewise, active immunization with neuromuscular junction proteins can cause myasthenic-like features in mice *(27)*. Although synaptic membrane proteins have been implicated in autoimmune encephalitis, studies of these diseases have largely been limited to passive transfer approaches *(3)*. Active immunization has the potential to improve our understanding of the evolution of the disease and act as a model for therapeutic interventions.

### Comparison to prior studies

In anti-NMDA receptor encephalitis, passive transfer approaches using patient-derived CSF/IgG *(12–14, 28)* provide compelling evidence for the involvement of NMDA receptor antibodies in pathogenesis *(13, 14, 20, 28)*. Tissue samples from patients with anti-NMDA receptor encephalitis also indicate that reactive gliosis and peripheral immune cell infiltrates are prominent pathological features *(12, 15–17)*. Intraventricular infusion of human NMDA receptor antibodies in mice decreases NMDA receptor density in the hippocampus *(14, 29)*, but does not recapitulate the gliosis and immune cell infiltrates, nor do these mice develop seizures or hyperactivity *(14, 29)*. These observations suggest that passive infusion of NMDA antibodies does not fully mimic the clinical syndrome. Our results show that active immunization of immune competent mice with NMDA receptor holoproteins induces a fulminant encephalitis that is strikingly similar to the human disease. Immunized mice show behavioral abnormalities, seizures, neuroinflammation, immune cell infiltrates and the presence of NMDA receptor specific antibodies. The intense labeling of hippocampus with IgG in our experiments also fits with the regional pattern in human cases of limbic encephalitis (Fig. S1).

### What causes disease symptoms?

The current hypothesis for the clinical phenotype is that antibody-mediated internalization of NMDA receptors leads to hypofunction at a network level. The available evidence convincingly demonstrates that antibodies from human cases cause internalization of NMDA receptors *in vitro* and reduce NMDA responses *(12, 13, 30)*. Thus therapies in use or proposed have been directed at removing antibodies, immunosuppression, or preventing receptor internalization *(3)*. The hypofunction hypothesis gains further support from the behavioral side-effects observed with use of NMDA receptors antagonists, but NMDA receptor hypofunction does not easily account for the presence of seizures or neuroinflammation. Our results with active immunization of mice show early and prominent neuroinflammation as well as histological evidence of neuronal loss. These features have also been observed in human cases *(12, 16, 17)*.

The extent to which these aspects contribute to the clinical signs and symptoms remains to be determined, but suggests a more complex pathogenesis than simply a reduction in NMDA receptor function.

### Role of T cells

Although prior studies of anti-NMDA receptor encephalitis have largely focused on the role of B cells and antibodies, the role of CD4^+^ T cells in autoimmune encephalitis is an area of growing interest *(31)*. T cells could promote neuroinflammation as well as potentiate B cell- and plasma cell-mediated antibody responses. Indeed, mice at three weeks post-immunization showed reactive gliosis, CD4^+^ T cell and galectin3^+^ macrophage infiltrates, whereas serum antibodies were still below detection by Western blots.. In human cases, CSF cytokine/chemokine profiles support a role for CD4^+^ T cell involvement *(32–35)*. For example, interleukin-17, a pro-inflammatory cytokine produced by Th17 CD4^+^ T cells, is prominent in the CSF of human cases and may perform the dual role of blood-brain barrier disruption and upregulation of IL-6, a pro-B cell and plasma cell cytokine *(32, 36, 37)*. The presence of CD4^+^ T cells in both early and later stages, and the occlusion of the disease in the TCRα^−^ mice lacking mature CD4^+^ or CD8^+^ cells support an important role for T cells in disease pathogenesis. Thus therapies designed to reduce T cell-mediated inflammation (e.g. blocking interleukin 17 signaling) as well as those aimed at reducing B cell activation are worthy of further investigation.

### Conformationally-restricted triggers

The trigger for an autoimmune reaction to NMDA receptors remains unclear. Immunization with peptide fragments used to produce NMDA receptor antibodies have not resulted in reports of clinical disease. In a recent study, mice immunized with NMDA receptor peptides did not show clinical symptoms, but the authors suggested that blood-brain barrier integrity may prevent circulating NMDA receptor antibodies from entering the CNS *(38)*. The immunogen in our case was the tetrameric GluN1/GluN2B receptor in a native-like heteromeric assembly *(39)*. These subunits had been altered to maximize protein stability, e.g. by removal of the intracellular C-termini *(39)*, and were capable of binding glutamate or glycine, which may also have improved protein stability. These factors may have played a role in the high incidence of disease in our mice, and may be relevant considerations for other membrane proteins implicated as causes of encephalitis. Our results strongly suggest that disease induction depends on conformationally-restricted epitopes. This idea is consistent with prior studies in HEK293 cells, which showed an assembled NMDA receptor was necessary for reactivity of antibodies from human cases *(4, 6)*. In human cases, the source of the intact NMDA receptors could be ectopic expression from a tumor, or in cellular debris following an insult causing neuronal cell loss. For example, there have been reports of anti-NMDA receptor encephalitis following viral infection *(7, 40)*. An association with viral infection was also suggested in the single reported case in Knut, a polar bear at the Berlin zoo *(41)*.

### Significance to human disease

Despites the prevalence of anti-NMDA receptor encephalitis, many experimental questions remain unanswered given the lack of a well-validated animal model that recapitulates the signs and symptoms to a high degree. The mouse model described here provides such a platform and has already provided several new insights. For example, use of conformationally-stabilized holoproteins appears to be a critical component of immunogenicity, and our results already indicate a complex pathogenesis involving B and T cells and a prominent inflammatory component. Thus the initial steps in disease induction, the roles of specific immune cells and potential new therapies can now be tested. Our approach also may prove useful as a template for other autoimmune disease involving CNS membrane proteins

## Methods

### Study Design

We examined the effect of active immunization with NMDA receptor native-like holoproteins on normal adult mice. The aim of the study was to investigate autoimmunity to NMDA receptors in the context of anti-NMDA receptor encephalitis. Littermate controls (liposome or saline) of both sexes were used for all interventions. Results from liposome and saline controls did not differ and were thus combined for statistical analysis of “control” in some experiments as indicated. Criteria were established in advance based on pilot studies for issues including data inclusion, outliers, selection of endpoints, and sample size (see statistics section). All analyses were blinded. Observations within each animal were averaged and the value for N replicates reflects the number of animals. Details for each experimental technique and analysis are included in the text and figure legends.

### Animals

We used the following mouse strains: C57BL/6J (Jax #000664), BALB/cJ (Jax #000651), MuMt^−^ (B6.129S2-lghm^tm1Cgn^/J; Jax #002288), and Tcrα^−^ (B6.129S2-Tcrα^tm1Mom^/J; Jax #002116). BALB/cJ mice were used only in pilot studies and all reported data are from C57BL/6J mice. Homozygous MuMt^−^ and Tcrα^−^ were obtained directly from Jax for immediate use in experiments and were not bred in-house. Genotyping of homozygous mutant MuMt^−^ and Tcrα^−^ breeder mice was conducted by Jax prior to shipping of offspring. MuMt^−^ and Tcrα^−^ were housed in the OHSU super barrier facility whereas other strains were housed in standard OHSU mouse facilities. All mice were housed under a 12hr light/dark cycle at 75°F, 60% humidity, with ad libitum access to food and water (Picolab Rodent Diet 20, no.5053). The OHSU Institutional Animal Care and Use Committee approved all procedures.

### NMDA receptor proteoliposome preparation and immunization

NMDA receptor holoprotein production and purification followed existing protocols *(39, 42)*. Subunits from X.laevis were used in these experiments, which show 92% amino acid identity (96% similarity) with rodent NMDA receptor subunits *(39, 42)*. The constructs were optimized for stability to allow purification of holoprotein complexes. HEK293 GnTI^−^ cells were transduced with P2 baculovirus for GluN1Δ2 and GluN2BΔ2. At 60hrs post-transduction cells were collected and membrane-bound NMDA receptor holoprotein was isolated and purified using affinity and size-exclusion chromatography methods. Purified holoprotein was then reconstituted into proteoliposome for immunization as described previously *(42)*. For immunization, the NMDA receptor proteoliposome pellet was resuspended in sterile saline. Adult mice (PND60) received a subcutaneous injection, proximal to the right inguinal lymph node, with proteoliposome (25μg total NMDA receptor protein/200μl saline), liposome, or saline, followed by a booster at two weeks. Mice were observed daily (2 min duration) in their home cage for overt signs of encephalitis. Observer was blinded to treatment condition using a 0-5 scoring scale: *hyperactivity* (0=normal exploratory movement, 1=continuous running); *circling* within the long body axis (absent=0, 1=present); *hunched back & lethargy* (0=absent, 1=present); *clinical seizure* (0=absent, 1=present); *death* (0=absent, 1=present), modified from a scoring scale used for a mouse model of HSV-1 encephalitis *(43, 44)*.

### Behavioral assessments

Behavioral assessment mice were performed at 5-6 weeks post-immunization in the OHSU behavioral research core. Fourteen male and 14 female mice, housed individually, from each treatment group (proteoliposome, liposome, saline) were tested. Mice were acclimated to the behavioral suite for three days prior to testing. Researchers were blinded to experimental conditions. Behavioral testing was conducted between 7am-12pm during the light-on period. The planned behavioral test battery included zero maze, nesting, open-field, novel-object recognition, and context-cued fear conditioning. However, the profound locomotor phenotype made results from novel-object recognition and context-cued fear condition difficult to interpret. Thus only open-field, nesting behavior and zero maze were used. *<u>Open Field:</u>* Mice freely exploring a 40 × 40 cm Plexiglas arena. Ethovision (Noldus, Netherlands) video tracking software was used to trace and quantify movement. Mice were acclimated to the arena for 5 min on day 1. On Days 2 and 3 mice were recorded for 10 min. We assessed total distance moved (cm) and percent time spent exploring the central region (interior 20 × 20 cm) of the arena *(45)*, *<u>Nesting Behavior:</u>* Assessment of nest building, a complex stereotyped behavior *(46)*. At time point zero two new pressed cotton nestlets were weighed, placed in the home cage of an individually housed mouse, and photographed. At 24 hours, a new photograph was taken and any untorn nesting material (pieces > 0.1 grams) was weighed and returned to the cage. A photograph was repeated at a 48 hours. Scoring of the nest was based on a 1-5 scale per Deacon et al (2006 from untouched nestlets (score 1) to a fully constructed nest with central cavity and surrounding wall (score 5). *<u>Zero Maze:</u>* The zero maze (Kinder Scientific, Poway, CA) consisted of four sections alternating between closed and open areas. Mice were placed in an open arm of the maze and allowed to explore for 10 min. Movement and location of the mice was tracked by an automated photo beam detection system. Outcome measures were total distance moved (cm) and percent time spent in open and closed areas of the maze *(45)*.

### Histology and immunohistochemistry

Brain tissue for histology and immunohistochemistry was prepared as follows: terminally anesthetized mice were transcardially perfused with 10 ml phosphate buffered saline (PBS) followed by 10ml of 4% paraformaldehyde (PFA) in PBS, and brains post-fixed in 4% PFA/PBS overnight. For hematoxylin & eosin (H&E) PFA fixed brains were transferred to 70% ethanol, paraffin embedded, and 6 μm microtome sections were mounted and stained. For Immunohistochemistry, floating vibratome sections (40-100 μm) were permeabilized, and nonspecific staining blocked in 0.4% Triton + 10% normal horse serum (NHS) in PBS for 2 hours at room temperature. Sections were incubated overnight (4°C) with primary antibody in 1.5% NHS-PBS, washed in PBS (3 × 15 min) prior to incubation with secondary antibodies (Invitrogen AlexaFluor, 1:300) in 1.5% NHS-PBS for 2hrs at room temperature. Sections were counterstained with DAPI (1 x15 min; Sigma # 28718-90-3, 1:20K in PBS) and mounted on slides using Fluoromount-G (SouthernBioTech # 01001). The following primary/secondary antibody pairings were used to label floating brain section: CD45R (BD Biosciences #550539, 1:25; Invitrogen #A21208, AlexaFluor488), CD4 (BD Biosciences #550280, 1:25; Invitrogen #A21208, AlexaFluor488), CD8 (BD Biosciences #550281, 1:25; Invitrogen #A21208, AlexaFluor488), CD20 (Santa Cruz Biotechnology #7735, 1:50; Invitrogen #A21447, AlexaFluor647), CD138 (R&D Systems #AF3190, 1:100; Invitrogen #A11057, AlexaFluor568), Galectin3 (R&D Systems #AF1197, 1:500; Invitrogen #A21447, AlexaFluor647), Iba1 (Wako Laboratory Chemicals #019-19741, 1:500; Invitrogen #A21206, AlexaFluor488), GFAP-Cy3 (Millipore Sigma #C9205, 1:500), GluN2A (Invitrogen #480031; Invitrogen #A11008 AlexaFluor488).

Immune cells (CD45^+^, CD4^+^, CD8^+^, CD20^+^, CD138^+^, Galectin3^+^) and inflammatory cells (Iba1, GFAP) were quantified in three representative brain sections spanning −0.9 mm to −2.3 mm from bregma *(47)* from six randomly chosen mice (3♀; 3♂) per group. The observer was blinded to experimental conditions. Images were captured using a Zeiss/Yokogawa CSU-X1 Spinning Disk microscope (10x 0.45 numerical aperture). A 4×4 tiled z-stack consisting of 26, 1.39 um, optical sections was collected. Exposure time and laser intensity settings were held constant during image collection. Fused max intensity projections were constructed using Zen Blue software (Zeiss) and Fiji-Image J software. Cells/μm^3^ reported reflect mean density values across sampled sections. Iba1 and GFAP immunofluorescence intensity measurements were plotted as arbitrary units of pixel intensity (Image J) using the following formula: fluorescence intensity = integrated density – (ROI area X mean background fluorescence).

### Serum collection, IgG purification and Western blots

For serum collection, blood from transcardial puncture was allowed to clot at room temperature for 15min, centrifuged (2000rpm) at 4°C for 15 min and supernatant aliquots stored at −20°C and 4°C. A Nab Protein A/G spin kit (ThermoFisher Sci. #89950) was used for IgG isolation.

IgG concentration was determined using the Bradford dye-binding method (Bio-Rad #5000002). Serum and IgG isolates were tested against the following purified NMDA receptor subunits with C-terminal deletions as indicated: *Xenopus laevis*: GluN1-3a, GluN1-3aΔATD *(39, 48)*, GluN2A (AAI70552.1) residues 1-834, GluN2B (NP_001104191.1) residues 1-839; rat: GluN1-1a (U08261) residues 1-847, GluN1-1b (U08263) residues 1-868, GluN2A (AF001423) residues 1-866, and GluN2B (NP_036706) residues 1-868. NMDA receptor subunit constructs were cloned into pEGBacMam for transfection of tsA201 adherent cells (HEK 293T, ATCC CRL-11268) *(49)*. All constructs had a green fluorescent protein (GFP) tag at the carboxyl terminus. Transfected tsA201 cells were solubilized in 200 μl of lysis buffer (150 mM NaCl, 20 mM Tris pH 8.0, 40 mM *n*-dodecyl-β-D-maltoside, 1 mM phenylmethylsulfonyl fluoride, 2 mg/ml leupeptin, 0.8 mM aprotinin, and 2 mM pepstatin A) for 1 hr. The extracts were centrifuged for 40 minutes at 40000 rpm at 4°C and the supernatants were collected, resolved by sodium dodecyl sulfate polyacrylamide gel electrophoresis using a 4-15% gradient gel and transferred to nitrocellulose. Anti-GFP antibody (Invitrogen, A11122), and serum samples were used at dilutions of 1:5000 and 1:100, respectively. The secondary antibodies (LI-COR IRDye 680, 926-68071 and IRDye 800CW, 926-32210) were used at a dilution of 1:10000.

### *in vitro* assays and immunocytochemistry

HEK293FT cells (ThermoFisher Sci. #R70007) were cultured on poly-D-lysine (0.2mg/ml PDL; Sigma # P6407) coated glass coverslips (1mm; Bellco Glass #1943-10012A) in DMEM medium (DMEM Gibco #11965-092; 10% HI-FBS Gibco #10082-139; 1% Pen/Strep ThermoFisher Sci #15140122; 1% MEM NEAA Gibco #11140-050; 1% Glutamax Gibco #35050-061). At 30% confluence, cells were transfected with rat wild-type GluN1-1a/pCI-neo and GluN2A/pCI-neo or GluN2B/pCI-neo constructs (a gift from Dr. Weinan Sun) via calcium-phosphate precipitation in a 1:1 ratio *(50)*. Transfected cells were incubated at 37°C. After four to six hours, the transfection solution was exchanged for fresh DMEM medium containing 10 μM (R)-3-(2-Carboxypiperazin-4yl)propyl-1-phosphonic acid (CPP; Abcam #ab120159), returned to the incubator for twelve hours followed by fixation in 4% PFA-PBS (10 min). For immunocytochemistry HEK cells were permeabilized and blocked for non-specific labeling as described above. The following primary/secondary antibody combinations were used for colabeling experiments: GluN1 (Abcam #ab9864R; 1:100; Invitrogen #A11008 AlexaFluor488, 1:400) co-incubated with purified IgG from liposome- or proteoliposome-treated mice (0.00625mg/ml; Invitrogen #A21236 AlexaFluor647, 1:400). A z-stack spanning a single cell layer was captured using a Zeiss/Yokogawa CSU-X1 Spinning Disk microscope (63x oil immersion 1.4 numerical aperture) with Zen Blue software (Zeiss) and Fiji-Image J software used for post-imaging processing.

For primary neuronal cultures, Hippocampi were dissected from C57BL/6 wildtype mice between post-natal days 0 and 2. Hippocampi were then dissociated with papain (Worthington LS003126; 200 U in 37°C dissection saline, 35 minutes) followed by trituration. Cultures were maintained in a tissue culture incubator at 37°C and 5% CO2 in media made with MEM (Gibco #11090-081), 5% heat inactivated fetal bovine serum (Gibco #10082-139), 2 mM Glutamax (Gibco #35050-061), 1ml/L Mito+ serum extender (Corning #CB50006) and 21 mM added glucose. Media was maintained in the tissue culture incubator and cultures had half the media volume changed every three days. All cultures were made in two cell plating stages. The first plating seeds the poly-D-lysine-coated coverslips (Neuvitro #GG-12-PDL) with glia (300,000-400,000 cells/35 mm dish) that were allowed to proliferate for at least two weeks. Neurons were then plated (100,000 - 200,000 cells/35 mm dish) in the second stage then grown on a glial substrate. For electrophysiological experiments, neurons were used after 7 - 10 days *in vitro*. Neurons were fixed in 100% methanol (−20°; 15min) for immunocytochemistry experiments. Staining protocols as described above were followed without additional permeabilization. NMDA receptor colabeling experiments were conducted at DIV14 and DIV21 using the following primary/secondary antibody combinations: GluN1 (Abcam #ab9864R, 1:100; Invitrogen #A11011 AlexaFluor568, 1:400), Map2 (Novus Biologicals #NB300-213, 1:500; Invitrogen #A11039 AlexaFluor488, 1:400), purified IgG from liposome- or proteoliposome-treated mice (0.00625mg/ml; Invitrogen #A21236 AlexaFluor647, 1:400). A Zeiss 880 laser scanning confocal microscope was used to acquire a z-stacked image (63x oil immersion 1.4 numerical aperture). Zen Blue software (Zeiss) and Fiji-Image J software used for image processing.

### Electrophysiology

For whole-cell voltage clamp recording, we used cultured mouse hippocampal neurons after 7 - 10 days *in vitro*. Recordings were done at room temperature, in external solution containing (in mM): 158 NaCl, 2.4 KCl, 10 HEPES, 10 D-glucose, 1 - 2 CaCl_2_, and 0.01 SR95531 (Abcam #ab120042). Solution pH was adjusted to 7.4 and osmolality adjusted to 320 mosmol/kg H_2_O. The pipette solution contained the following (in mM): 140 cesium methanesulfonate, 4 NaCl, 0.5 CaCl_2_, 5 EGTA, 10 HEPES, 4 magnesium-ATP, 0.3 sodium-ATP (pH 7.4; 320 mosmol/kg H_2_O). Neurons were continuously perfused through gravity-fed flow pipes which were placed within 100 micrometers of the recorded neuron during recordings. Cells were voltage clamped at −70 mV. Recording electrodes had resistances of 1.5 - 2.5 MOhm. The recording series resistance was always <8 mOhm and was compensated by at least 80% using the amplifier circuitry. Tetrodotoxin (1 μM) and NBQX (2.5 μM) were used in NMDA application experiments and drugs were applied via custom flow pipes that were translated with a computer-controlled piezo-electric bimorph. Computer-controlled solenoid-gated flow pipes were used for delivery of NMDA. In a different set of experiments, neurons were incubated with either serum from proteoliposome or liposome-treated mice for 24 hours and spontaneous excitatory postsynaptic currents (sEPSCs) were recorded at a holding potential of −70 mV with or without D-AP5 (50μM; Abcam #ab120003). Axopatch 1C amplifiers and AxoGraph acquisition software (Axograph X) were used for data acquisition. Data were low-pass filtered at 2.5-5 kHz and acquired at 5-10 kHz.

### Statistics

Sample size were determined based on prior experiments of this type with an effect size of 20% and power = 0.8. Tests of normality were used to determine the appropriate test. Multiple comparisons used 1- or 2-way ANOVA or nonparametric ANOVA as indicated for each experiment. Exact p values are provided and both the number of animals and the number of observations are indicated as appropriate. Data unless other indicated are plotted as mean ± SEM. All statistical analyses were completed using Prism 7 software (GraphPad).

## Acknowledgments

This work was supported by NS080979 and Ellison Medical Foundation (GLW), the NINDS imaging core facility (P30NS061800) and by NS038631 (EG). E.G. is an Investigator of the Howard Hughes Medical Institute. We thank the OHSU histology core for processing of samples, Randy Woltjer for advice concerning interpretation of the histology, Weinan Sun for wild-type NMDA receptor subunit constructs, and Farzad Jalali-Yazdi for NMDA receptor amino acid sequence alignments.

## List of Supplementary Materials

**Fig S1.** IgG labeling in the CNS of proteoliposome treated-mice.

**Fig S1.** Neuroinflammation and immune cell infiltrate at 3 weeks post-immunization.

**Fig S1.** Neuroinflammation and immune cell infiltrate at 3 weeks post-immunization.

**Fig S1.** B cells are required to generate disease state in proteoliposome treated mice.

**Movie S1-S5:** Home cage video illustrating behavioral phenotype of proteoliposome-treated mice.

